# Comparison of bacterial communities from the surface and adjacent bottom layers water of Billings reservoir

**DOI:** 10.1101/2021.06.03.447016

**Authors:** Marta Angela Marcondes, Andrezza Nascimento, Rodrigo Pessôa, Jefferson Russo Victor, Alberto José da Silva Duarte, Sabri Saeed Sanabani

## Abstract

Here, we describe the microbial diversity and physicochemical properties in freshwater samples from the surface and bottom layer of Billings reservoir in São Paulo state, Brazil. Twenty-two matched samples were characterized using the 16S rRNA gene Illumina MiSeq platform. Taxonomical composition revealed an abundance of *Cyanobacteria* phyla, followed by *Proteobacteria*, with 1903 and 2689 known bacterial genera in the surface and deep-water layers, respectively. Shannon diversity index ranging from 2.3 - 5.39 and 4.04 - 6.86 in the surface and bottom layer, respectively. Among the 120 pathogenic genera identified, *Flavobacterium* was the most predominant genus. Temperature and phosphorus concentration were the most influential factors in shaping the microbial communities of both layers. Predictive functional analysis suggests that the reservoir is enriched in motility genes involved in the flagellar assembly. The overall results present new information on the significantly altered diversity composition of the bacterial community detected in Billings freshwater reservoir.

**IMPORTANCE:** In this study, we investigated the bacterial distribution, community composition, potential metabolic activity, potentially pathogenic bacteria, and toxin genes of *Cyanobacteria* in the bottom layers and surface along Billings reservoir in the southeast of Brazi. Our results provide essential information about the pattern of bacterioplankton communities’ variation inhabiting the Billings reservoir and the combination of environmental that shaped their structure. These results may help pave the way for future studies devoted to control and improve the water quality in the Billings reservoir, which is facing rapid urban development and urbanization.

## 1. INTRODUCTION

The Billings reservoir in the State of São Paulo, Brazil, is the largest freshwater storage aquatic body in the São Paulo Metropolitan Region, which covers 127 km2, have a total volume of 1228.7 × 106 m3 and a maximum depth of 18 meters (1, 2). The reservoir has multiple-use, including hydropower generation, water supply for 4.5 million people, and industries, irrigation, fishery, and flood control. The basin has a narrow central body which measures over 20 km and to which eight branches, namely Rio Grande, Rio Pequeno, Capivari, Pedra Branca, Taquacetuba, Bororé, Cocaia, and Alvarenga rivers, converge (3–5). The reservoir has undergone considerable changes in its properties since the 1940s. At that time, part of the polluted water from the Tiete River (São Paulo city) was allowed to flow into the reservoir, believing that it would raise the water level to generate electric power (6). This process, besides the pressure of uncontrolled urban growth, significantly contributed to considerable anthropogenic eutrophication and algal bloom of the Billings reservoir (7).

Reliable access to clean and affordable water is one of the most basic humanitarian goals and is a major global challenge for the 21st century (8, 9). Water pollution is caused by the discharge of harmful domestic and industrial wastes into surface water bodies like rivers, dams, reservoirs, lakes, and canals due to the lack of or inefficient wastewater treatment plants (10–12).

Reservoirs are critical artificial aquatic bodies of many drinking water supply systems; they can maintain an equilibrium of storing or releasing water, play a central role in the biogeochemical cycling, energy flows, and the recycling of nutrients. The functioning of these cycling is mainly maintained by the inhabiting microorganisms. Despite their potential importance, the structure and functioning of the microbial communities in these ecosystems have received relatively less attention compared to other freshwater bodies, such as natural lakes and rivers (13). There are very few studies on bacterial community structures and compositions of the surface waters in Brazil (14, 15). Thus, this study aimed to (i) to explore and compare the microbial communities in surface and bottom layer water along the Billings reservoir using the 16 S rRNA gene-based Illumina MiSeq sequencing, (ii) evaluate the presence of potential pathogens in these water samples and (iii) explore the predicted functional profiles of the obtained microbial communities in the basin to determine their role in the ecosystem.

## 2. MATERIALS AND METHODS

### 2.1 Study Sites and Sample Collection

The study area covered the entire 127 km2 of the Billings reservoir, which is located west of the city of Sao Paulo at 230 47’S, 460 40’W, W, an altitude of 746 m a.s.l. Surface and bottom layer water samples were collected in March 2019 from 30 locations (about 17 km apart; **Figure 1**) in triplicate using a Van Dorn sampler as previously described (11). Water temperature (Temp), pH, dissolved oxygen (DO), specific conductance (SPC), pH, and Chloride ion were evaluated on-site from each sample using a handheld multi-parameter water quality sonde (YSI Inc./Xylem Inc.) Other variables including turbidity, nitrate (NO3-), sulfate (SO4–2), orthophosphate (PO43−), phosphorus (P), and ammonia nitrogen (NH4+-N) were determined according to the Brazilian standard issued by Environment National Council (CONAMA resolution 357/2005). Table 1–4 provide a list of sample sites, their physical description, and representative physiochemical data.

**Table 1.** Physiochemical characteristics of water samples from the bottom layer of Billings reservoir.

| **Sample ID** | **DO\* mg/l** | **Temp** | **pH** | **Average Water Depth (Km)** | **Turbidity (NTU)** |
| --- | --- | --- | --- | --- | --- |
| B1.1F | 3.7 | 20.6 | 7.43 | 9.5 | 4.13 |
| B2.1F | 6.4 | 20.8 | 8.25 | 5.5 | 3.03 |
| B3.1F | 5.1 | 19.8 | 7.21 | 3.2 | 3.86 |
| B4.1F | 4 | 18.8 | 7 | 2.1 | 1.16 |
| B5.1F | 7.1 | 21.1 | 8.23 | 6.9 | 2.99 |
| B6.1F | 7.1 | 21.7 | 8.28 | 12.8 | 4.38 |
| B7.1F | 5.6 | 21.2 | 8.07 | 12.4 | 6.28 |
| B8.1F | 4.6 | 21.1 | 7.91 | 10.1 | 8.08 |
| B9.1F | 4.5 | 20.9 | 7.18 | 9 | 9.81 |
| B10.1F | 4.8 | 21.2 | 7.64 | 7.6 | 13.55 |
| B11.1F | 3.5 | 21.2 | 7.55 | 5.9 | 10.3 |
| B12.1F | 3.5 | 20.9 | 7.41 | 8.1 | 6.72 |
| B13.1F | 4.9 | 20.8 | 7.66 | 10.4 | 10.27 |
| B14.1F | 8.3 | 21 | 8.27 | 3.9 | 31.9 |
| B15.1F | 4.3 | 20.8 | 7.53 | 13.6 | 6.09 |
| B16.1F | 8.4 | 20.6 | 8.51 | 10.1 | 11.5 |
| B17.1F | 6.3 | 20.5 | 7.8 | 6.9 | 16.5 |
| B18.1F | 6 | 20.3 | 7.63 | 8.9 | 13.1 |
| B19.1F | 5.9 | 20.5 | 7.64 | 6.1 | 9.93 |
| B20.1F | 5.5 | 20.4 | 7.51 | 4.5 | 9.87 |
| B21.1F | 6.6 | 20.5 | 7.73 | 7 | 8.36 |
| B22.1F | 7.8 | 20.5 | 8.38 | 5.4 | 10.2 |
| B23.1F | 6.9 | 20.9 | 8.1 | 5.8 | 8.16 |
| B24.1F | 7.1 | 20.7 | 7.96 | 5.5 | 6.79 |
| B25.1F | 6.6 | 20.9 | 7.92 | 6.9 | 6.15 |
| B26.1F | 7.6 | 21.3 | 8 | 6.8 | 7.78 |
| B27.1F | 8.3 | 20.5 | 7.81 | 3.2 | 3.4 |
| B28.1F | 7.6 | 20.5 | 7.9 | 7.1 | 7.19 |
| B29.1F | 8.2 | 20.6 | 8.63 | 8.7 | 7.72 |
| B30.1F | 7.7 | 20.5 | 8 | 10.2 | 7.6 |

\*Dissolved Oxygen

| Sulfate mg/L | Phosphorus mg/L | Ammonia mg/L | Nitrate mg/L |
| --- | --- | --- | --- |
| 0.4 | 0.12 | 2.5 | 1.1 |
| 0.5 | 0.2 | 2.3 | 1.2 |
| 0.4 | 0.23 | 2.8 | 1.5 |
| 0.1 | 0.14 | 2.4 | 1.2 |
| 0.6 | 0.08 | 2.3 | 0.05 |
| 0.4 | 0.1 | 3.2 | 0.09 |
| 0.5 | 0.09 | 2.8 | 1.2 |
| 0.2 | 0.08 | 2.2 | 1.1 |
| 0.4 | 0.043 | 2.4 | 0.9 |
| 0.5 | 0.1 | 3.7 | 0.8 |
| 0.2 | 0.04 | 3.1 | 1.1 |
| 0.3 | 0.04 | 2.8 | 0.8 |
| 0.5 | 0.03 | 3.1 | 1.2 |
| 0.3 | 0.02 | 2.5 | 0.9 |
| 0.5 | 0.06 | 2.7 | 1.2 |
| 0.4 | 1.2 | 2.3 | 0.09 |
| 0.4 | 1.1 | 2.6 | 1.1 |
| 0.4 | 0.08 | 2.7 | 0.09 |
| 0.8 | 1.1 | 2.6 | 1 |
| 0.6 | 2.1 | 2.4 | 2.7 |
| 0.3 | 0.9 | 2.8 | 1.1 |
| 0.6 | 1.2 | 2.7 | 1.6 |
| 0.5 | 1.3 | 2.3 | 1.8 |
| 0.4 | 0.9 | 3 | 2.1 |
| 0.5 | 1.2 | 3.5 | 1.9 |
| 0.5 | 1.9 | 2.9 | 2.3 |
| 0.6 | 2.1 | 3.1 | 1.9 |
| 0.4 | 1.9 | 2.9 | 1.6 |
| 0.5 | 2.1 | 2.4 | 1.1 |
| 0.5 | 0.9 | 2.3 | 1.3 |

**1.**
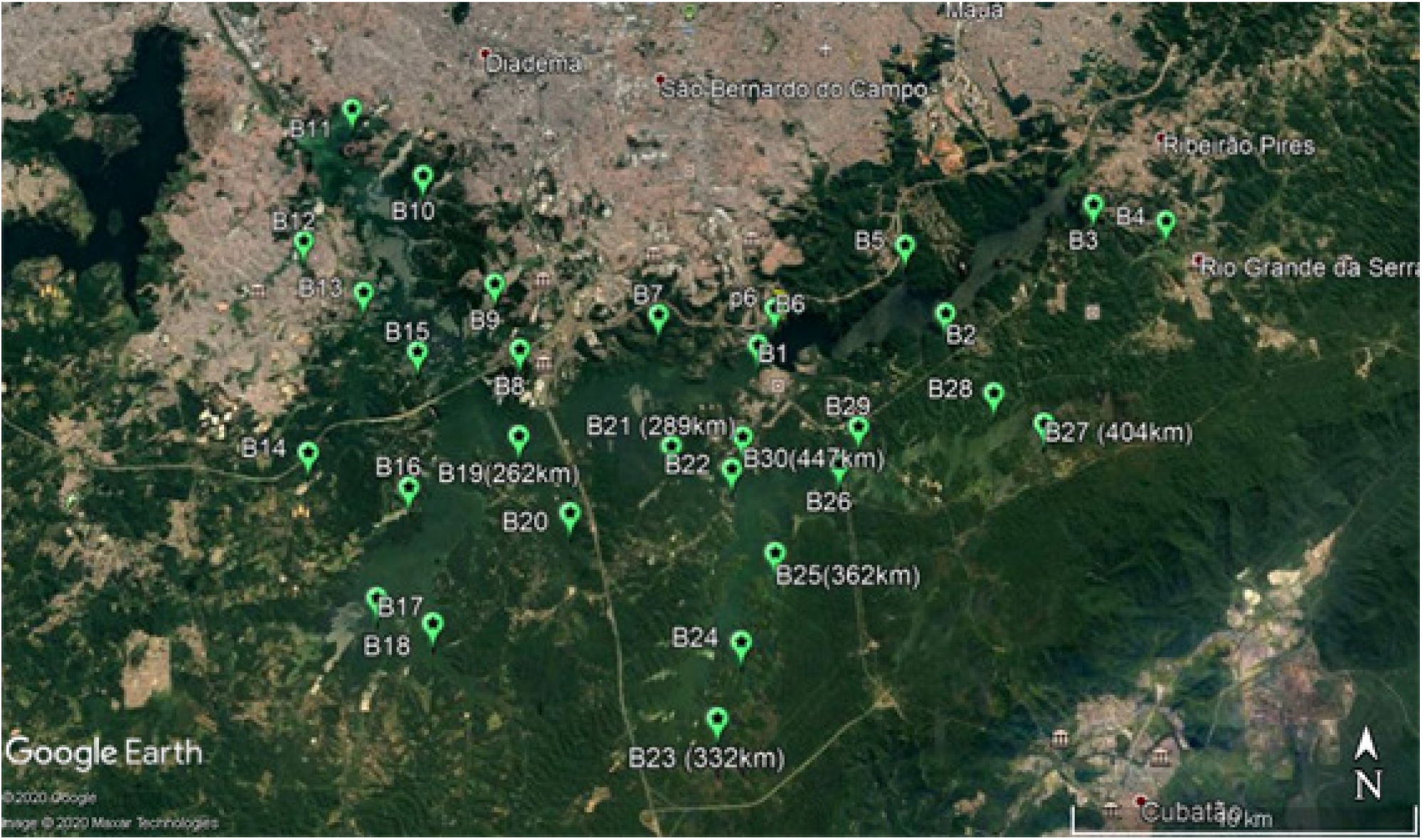
Map showing sampling site locations in the Billings reservoir in São Paulo. Locations are indicated by the green Global Positioning System (GPS) symbol. Map was obtained using the Google Earth mapping engine (https://www.google.com/earth/).

### 2.2 DNA Isolation, Gene Amplification, and library preparation

The total genomic DNA from each point was extracted using the PowerSoil DNA kit (MO BIO Laboratories™: Carlsbad, CA, USA) as per the manufacturer’s instructions. To minimize the potential bias during DNA extraction, each sample was extracted in duplicate and then pooled to quantify their DNA yield with *a Qubit*® fluorometer (Invitrogen, USA). The extracted DNA from each sample was subjected to PCR amplification of the V3-V4 variable region of the 16S rRNA gene using the previously published primers Bakt_341F/Bakt_805R (16) and conditions previously described by our group (17, 18). Library preparation and massively parallel sequencing (MPS) were performed as previously reported (11, 17, 18).

### 2.3 Detection of toxin-producing cyanobacterial genes

Three regions of the microcystin synthetase (*mcy*) gene cluster was selected to search for potential microcystins (MCs) producers in Billings samples. Representative samples (n = 15) with cyanobacteria in > 40% of their bacterial communities were selected for assay. A different set of specific published primers designed to detect mcyA (mcyA-Cd_1F; 5’-AAA ATT AAA AGC CGT ATC AAA-’3 and mcyA-Cd_1R; 5’-AAA AGT GTT TTA TTA GCG GCT CAT-’3) (19), mcyD (mcyDF; 5’-GAT CCG ATT GAA TTA GAA AG-’3 and mcyDR; 5’-GTA TTC CCC AAG ATT GCC-’3), and mcyE gene (mcyE-F2; 5’-GAA ATT TGT GTA GAA GGT GC-’3 and mcyE-R4; 5’-AAT TCT AAA GCC CAA AGA CG-’3) (20). The amplification for all fragments contained 50–100 ng DNA template, 2 mM MgCl2, 0.1 mM dNTPs, 0.5 μM of each primer, and 2.5 U high-fidelity *Taq* platinum DNA polymerase (Invitrogen, Carlsbad, CA) in a MgSO_4_ reaction buffer. After an initial denaturation of 5 min at 94°C, 35 cycles of 30s at 94°C, 30s at 55°C, 60s at 72°C and a final extension at 72°C for 5 min were performed. Each PCR included a known cyanobacterial DNA positive control, and an interspersed no DNA template negative control. The amplified product was electrophoresed through 1% (wt/vol) agarose gels containing 0.5 × Tris Borate EDTA, followed by ethidium bromide staining.

### 2.4 Bioinformatics and Statistical Analysis

Base-calling and data quality were initially assessed on the MiSeq instrument using RTA v1.18.54, and MiSeq Reporter v2.6.2.3 software (Illumina Inc., CA). The sequences were analyzed by a pipeline of the 16s Microbiome Taxonomic Profiling of the EzBioCloud (https://www.ezbiocloud.net/) application and the EzBioCloud Database Update 2019.04.09. Briefly, the analysis includes the quality-controlled 16S reads, taxonomic assignment, and the estimation of the functional profiles of the microbiome identified using 16S rRNA sequencing by the PICRUSt algorithm. The predicted profiles were categorized into clusters of Kyoto Encyclopedia of Genes and Genomes (KEGG) orthology and KEGG pathways. The differences in the surface and bottom layer characteristics were investigated between the groups using the t-test. Bacteria richness was measured by Chao1 and the operational taxonomic units (OTUs) number detected in the microbiome taxonomic profile (MTP) index. The ACE, Chao1, and Jackknife α-diversity indices were used to calculate the bacterial richness and the Shannon, Simpson function, and NPShannon indices to estimate evenness in each group using the Wilcoxon rank-sum test. Beta diversity was computed with Jansen-Shannon distances based on the profiles of taxonomic abundance. The statistical significances of β-diversity were computed using the permutational multivariate analysis of variance (PERMANOVA). The enrichment in the assigned taxonomic and functional profiles of the two groups were defined by the linear discriminant analysis (LDA) of the effect size (LEfSe) algorithm. The correlations between bacterial community diversity and water properties were assessed using principal component analysis (PCA).

For the detection of bacterial pathogens, we considered any bacteria potentially pathogenic if at least one species with a minimum abundance of 10 strains of any genus was categorized as biosafety level 2 or 3 by the American Biological Safety Association (https://my.absa.org).

### 2.5 Sequence data availability

All sequence data described here are available in the online Zenodo repository: https://doi.org/10.5281/zenodo.4751698

## 3. RESULTS

### 3.1 Physicochemical characteristics water samples

Samples were collected in June 2019 where the temperature was above 20^0^C the season was slightly dry. The water depth of the study area from the reservoir ranged from 1.16 m to 13.6 m, depending on the locations of collected samples. **Table 1 & 2** depict the physicochemical properties of the surface and bottom layer water samples from Billings’s reservoir at 30 different sites. Temperature and DO ranged from 18.8 °C to 22.1 °C and 3.5 to 9.5, respectively. Both parameters are significantly higher (p <0.5) in surface water when compared to the bottom layer (Average temperature: 21.1 °C versus 20.7 °C, Averaged DO: 9.5 mg/L versus 8.4 mg/L), while the phosphorus concentrations showed the opposite patterns. To test possible associations between the bacterial communities and the physicochemical parameters, PCA was performed (**Figure 2 & 3**) and the results revealed that the bacterial communities in the bottom layer were most correlated to DO, depth and temperature. The same analysis in samples from surface water showed the most positive correlation with ammonia concentration and temperature. The first PCA axis explained 29% and 26.4% of the variation of bacterial communities, and the second explained 24.1 and 18.1% of the surface and bottom layer water, respectively.

**Table 2.** Physiochemical characteristics of water samples from the surface of Billings reservoir.

| Sample ID | DO* mg/l | Temp | pH | Average Water Depth (Km) | Turbidity (NTU) | Sulfate mg/L |
| --- | --- | --- | --- | --- | --- | --- |
| B1.1S | 4 | 20.5 | 7.28 | 9.5 | 3.05 | 0.4 |
| B2.1S | 7.1 | 21.2 | 8.27 | 5.5 | 3.08 | 0.5 |
| B3.1S | 5.7 | 21.2 | 7.45 | 3.2 | 2.5 | 0.4 |
| B4.1S | 5.5 | 22.1 | 6.97 | 2.1 | 1.86 | 0.6 |
| B5.1S | 7.4 | 21.5 | 8.13 | 6.9 | 2.51 | 0.4 |
| B6.1S | 8.3 | 21.6 | 8.65 | 12.8 | 4.28 | 0.3 |
| B7.1S | 7.5 | 21.2 | 8.15 | 12.4 | 6.91 | 0.3 |
| B8.1S | 7.3 | 21.1 | 7.9 | 10.1 | 15 | 0.4 |
| B9.1S | 7.2 | 21.2 | 7.58 | 9 | 11.5 | 0.7 |
| B10.1S | 7.9 | 21.5 | 7.93 | 7.6 | 18.53 | 0.3 |
| B11.1S | 5.8 | 21.5 | 7.57 | 5.9 | 12 | 0.6 |
| B12.1S | 4.3 | 21.2 | 7.52 | 8.1 | 14.1 | 0.5 |
| B13.1S | 6.7 | 20.8 | 7.8 | 10.4 | 10 | 0.5 |
| B14.1S | 9.5 | 20.9 | 8.46 | 3.9 | 12 | 0.54 |
| B15.1S | 4.7 | 20.9 | 7.5 | 13.6 | 6.22 | 0.56 |
| B16.1S | 9.2 | 21 | 8.63 | 10.1 | 12.9 | 0.4 |
| B17.1S | 7.9 | 20.6 | 7.73 | 6.9 | 11.8 | 0.6 |
| B18.1S | 6.5 | 20.5 | 7.47 | 8.9 | 18 | 0.4 |
| B19.1S | 6.3 | 20.5 | 7.56 | 6.1 | 7.82 | 0.5 |
| B20.1S | 6.5 | 20.9 | 7.3 | 4.5 | 10.5 | 0.3 |
| B21.1S | 7.7 | 20.9 | 7.58 | 7 | 12 | 0.5 |
| B22.1S | 9.4 | 20.6 | 8.29 | 5.4 | 11 | 0.5 |
| B23.1S | 9 | 21 | 7.28 | 5.8 | 6.5 | 0.4 |
| B24.1S | 7.9 | 21.3 | 8 | 5.5 | 8.07 | 0.5 |
| B25.1S | 5.8 | 21.8 | 7.61 | 6.9 | 5.05 | 0.5 |
| B26.1S | 7.6 | 21.6 | 8 | 6.8 | 6.52 | 0.4 |
| B27.1S | 9.2 | 20.8 | 7.4 | 3.2 | 5.31 | 0.3 |
| B28.1S | 9.3 | 20.6 | 7.9 | 7.1 | 4.79 | 0.4 |
| B29.1S | 9.3 | 21.6 | 8.86 | 8.7 | 5.28 | 0.5 |
| B30.1S | 8.1 | 20.9 | 7.78 | 10.2 | 8.3 | 0.4 |

\*Dissolved Oxygen

| Phosphorus mg/L | Ammonia mg/L | Nitrate mg/L |
| --- | --- | --- |
| 0.08 | 2.4 | 0.9 |
| 0.21 | 2.8 | 1.1 |
| 0.06 | 2.8 | 2.2 |
| 0.02 | 2.7 | 7.4 |
| 0.17 | 2.7 | 0.9 |
| 0.32 | 2.2 | 1.2 |
| 0.06 | 2.6 | 0.9 |
| 0.27 | 2.5 | 1.2 |
| 0.3 | 2.5 | 1 |
| 0.84 | 2.9 | 2.2 |
| 0.12 | 3 | 1.8 |
| 0.02 | 2.6 | 1.4 |
| 0.14 | 2.7 | 1.9 |
| 0.12 | 3.3 | 1.8 |
| 0.03 | 2.7 | 1.7 |
| 0.11 | 2.8 | 1.3 |
| 0.09 | 2.7 | 2.3 |
| 0.15 | 2.7 | 2.65 |
| 0.09 | 2.1 | 1.3 |
| 0.11 | 2.6 | 2.9 |
| 0.16 | 2.8 | 2.2 |
| 0.19 | 2.1 | 1.1 |
| 0.15 | 2.4 | 1.8 |
| 0.04 | 2.9 | 1.02 |
| 0.05 | 3.4 | 1.98 |
| 0.11 | 3.2 | 0.9 |
| 0.08 | 2.4 | 0.8 |
| 0.11 | 2.8 | 0.9 |
| 0.08 | 2.9 | 1.2 |
| 0.19 | 2.3 | 2.8 |

**2.**
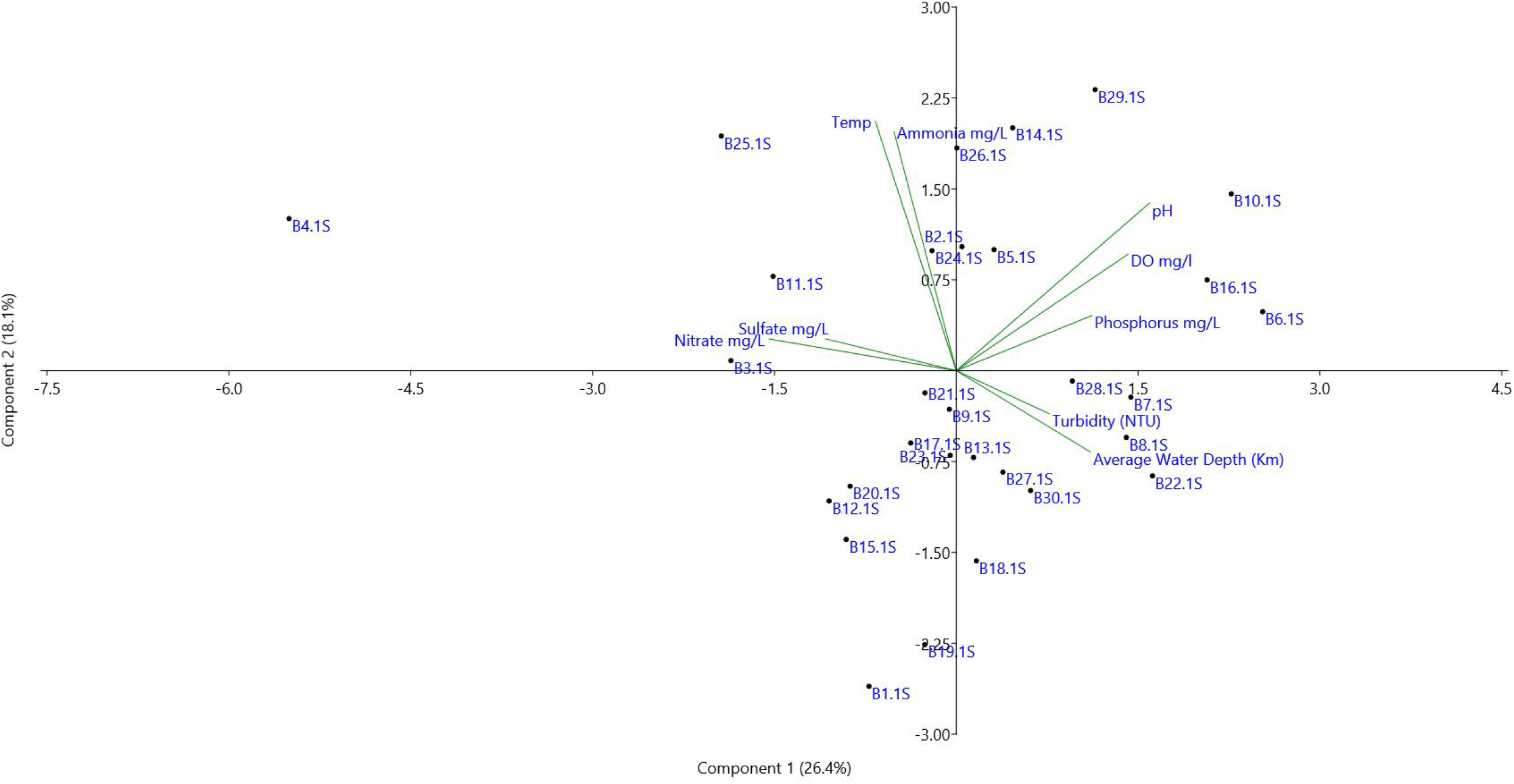
Principal component analysis (PCA) biplot of bacterial communities and surface water properties.

**3.**
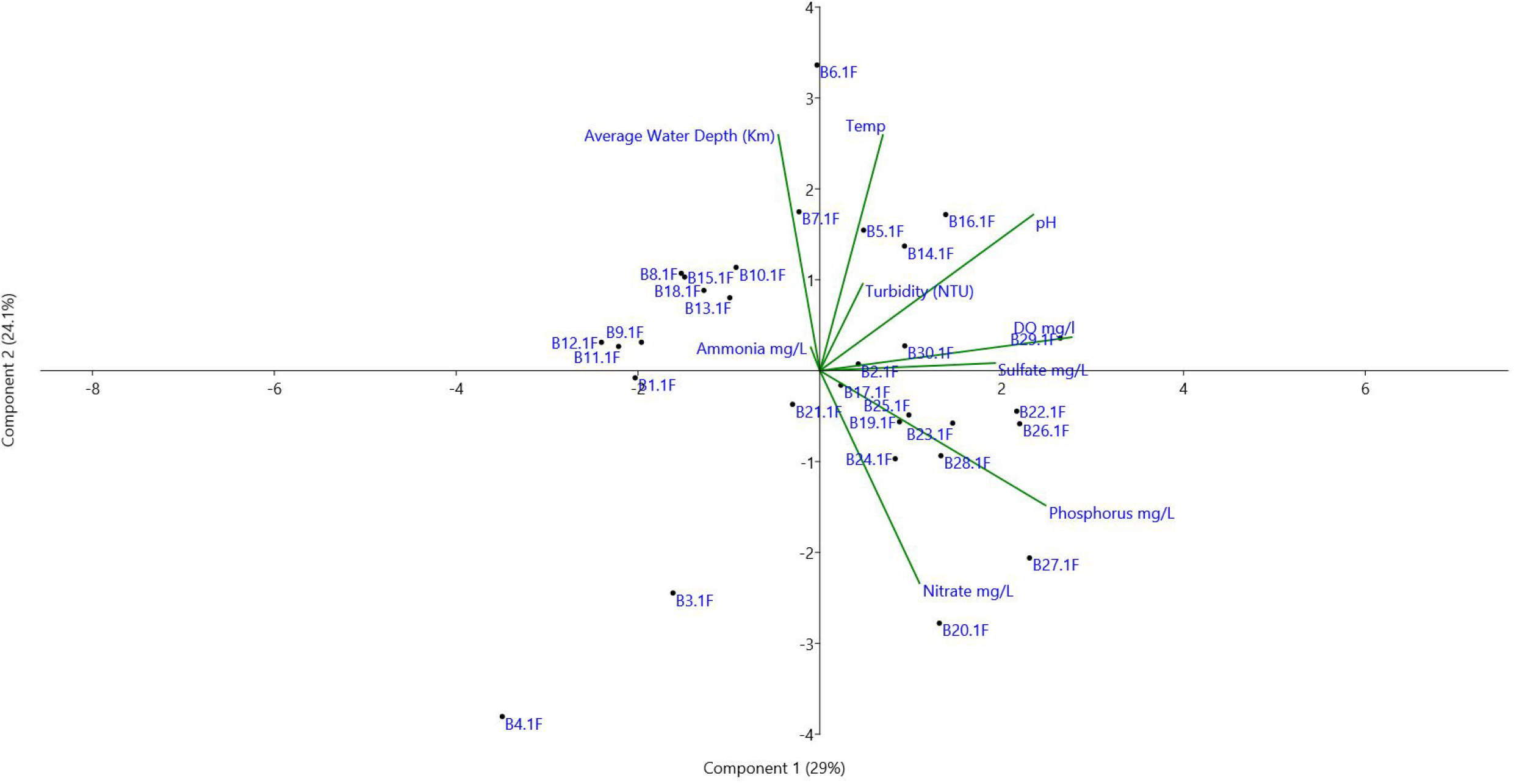
Principal component analysis (PCA) biplot of bacterial communities and bottom layer water properties.

### 3.2 Bacterial community structure

A total of 22 matched water samples from the surface and bottom layer were successfully amplified, sequenced, and submitted for further analysis. To minimize computational time, a random of 100,000 reads from each sample were selected, cleaned, and analyzed by the EzBioCloud tool. This resulted in 1,468,033 (Min: 15,766 in B7; Max: 87,913 in B21) and 1,861,126 (Min: 35,189 in B22F; Max: 96,434 in B12F) valid reads in surface and bottom layer water, respectively. The good’s coverage estimator of the OTUs in the surface and bottom layer samples ranged from 99.38 to 99.95% and 99.05 to 99.84, respectively (**Table 3 & 4**), showing that the diversity of bacterial populations in both groups was sufficiently covered by the generated sequences. The median Shannon’s diversity index showed no significant difference between the two layers (P >0.3). Both in surface and bottom layer water, the Shannon index of B04 was the highest at 5.39 and 6.19, respectively, indicating abundant community diversity. Of note, the only average of approximately 52% of the municipalities’ sewage in this area of the reservoir receives any kind of treatment, and another 48%, which could be treated, is not. Thus, the high community bacterial diversity in these areas possibly indicates that a substantial portion of the untreated waste is dumped into the reservoir. Our results also showed that the microbiota of the bottom layer had significantly higher phylogenetic diversity indices than those of the surface layer (p < 0.005) (**Figure 4**).

**Table 3.** The number of raw and valid reads sequenced for each matched sample from the surface water, number of species and OTUs found, and subsequent alpha diversity measures.

| MTP name | Valid reads | Valid reads | OTUs | ACE | CHAO | Jackknife | NPSannon |
| --- | --- | --- | --- | --- | --- | --- | --- |
| B4F | 87,428 | 83,79 | 2116 | 2184.78 | 2143.90 | 2281.00 | 5.43 |
| B5F | 51,832 | 84,101 | 1550 | 1618.55 | 1569.69 | 1688.00 | 5.33 |
| B7F | 15,766 | 92,44 | 798 | 851.92 | 819.60 | 896.00 | 5.14 |
| B8F | 65,835 | 91,036 | 1990 | 2127.32 | 2043.62 | 2236.00 | 4.85 |
| B9F | 84,508 | 76,82 | 433 | 457.00 | 445.58 | 473.00 | 2.38 |
| B10F | 45,319 | 60,821 | 305 | 358.06 | 337.18 | 365.00 | 2.84 |
| B11F | 64,536 | 93,392 | 1943 | 2112.04 | 2014.37 | 2229.00 | 4.35 |
| B12F | 81,047 | 96,434 | 1777 | 1919.64 | 1838.63 | 2021.00 | 3.65 |
| B13F | 72,29 | 95,249 | 1531 | 1673.73 | 1603.15 | 1761.00 | 4.01 |
| B14F | 70,107 | 91,409 | 1673 | 1810.66 | 1726.28 | 1902.00 | 4.82 |
| B15F | 72,382 | 92,202 | 1730 | 1848.16 | 1778.59 | 1941.00 | 4.03 |
| B16F | 68,314 | 95,132 | 1635 | 1789.01 | 1698.62 | 1879.00 | 4.56 |
| B18F | 65,941 | 92,605 | 1926 | 2054.21 | 1967.21 | 2156.00 | 5.22 |
| B19F | 67,777 | 92,066 | 2003 | 2103.75 | 2031.51 | 2203.00 | 5.01 |
| B20F | 62,497 | 86,364 | 2085 | 2232.55 | 2132.10 | 2347.00 | 5.20 |
| B21F | 87,913 | 88,101 | 1660 | 1758.23 | 1704.39 | 1841.00 | 5.04 |
| B22F | 68,781 | 35,189 | 1735 | 1902.48 | 1813.24 | 2009.00 | 4.88 |
| B23F | 74,19 | 77,192 | 1825 | 1927.49 | 1855.23 | 2015.00 | 4.78 |
| B24F | 57,108 | 90,412 | 1353 | 1404.05 | 1371.17 | 1464.00 | 4.77 |
| B25F | 67,702 | 90,585 | 2072 | 2176.56 | 2102.82 | 2282.00 | 5.08 |
| B29F | 69,843 | 66,79 | 2150 | 2282.51 | 2194.64 | 2407.00 | 5.22 |
| B30F | 66,917 | 88,996 | 1526 | 1586.43 | 1547.79 | 1654.00 | 5.01 |

ber of species and OTUs found, and subsequent alpha diversity measures.
| **Shannon** | **Simpson** | **Phylogenetic Diversity** | **Good's coverage of library(%)** |
| --- | --- | --- | --- |
| 5.39 | 0.01 | 2231 | 99.81 |
| 5.29 | 0.01 | 1655 | 99.73 |
| 5.08 | 0.02 | 1175 | 99.38 |
| 4.80 | 0.07 | 2304 | 99.63 |
| 2.38 | 0.40 | 808 | 99.95 |
| 2.83 | 0.18 | 685 | 99.87 |
| 4.30 | 0.10 | 2320 | 99.56 |
| 3.61 | 0.22 | 2264 | 99.70 |
| 3.97 | 0.15 | 1874 | 99.68 |
| 4.78 | 0.04 | 2020 | 99.67 |
| 3.99 | 0.17 | 2058 | 99.71 |
| 4.52 | 0.06 | 2024 | 99.64 |
| 5.17 | 0.02 | 2258 | 99.65 |
| 4.96 | 0.04 | 2308 | 99.70 |
| 5.14 | 0.02 | 2377 | 99.58 |
| 5.01 | 0.02 | 2235 | 99.79 |
| 4.85 | 0.03 | 2102 | 99.60 |
| 4.74 | 0.04 | 2120 | 99.74 |
| 4.74 | 0.03 | 1844 | 99.81 |
| 5.02 | 0.02 | 2213 | 99.69 |
| 5.17 | 0.02 | 2307 | 99.63 |
| 4.98 | 0.02 | 1932 | 99.81 |

**Table 4.** The number of raw and valid reads sequenced for each matched sample from the bottom layer water, number of species and OTUs found, and subseque

| **Sample name** | **OTUs** | **ACE** | **CHAO** | **Jackknife** | **NPSannon** | **Shannon** | **Simpson** |
| --- | --- | --- | --- | --- | --- | --- | --- |
| B4F | 5142 | 5581.14 | 5323.41 | 5940.00 | 6.30 | 6.19 | 0.01 |
| B5F | 1630 | 1788.52 | 1717.55 | 1877.00 | 5.25 | 5.22 | 0.02 |
| B7F | 1398 | 1477.34 | 1432.59 | 1544.00 | 5.06 | 5.04 | 0.02 |
| B8F | 1700 | 1812.65 | 1758.67 | 1902.00 | 4.70 | 4.67 | 0.07 |
| B9F | 1529 | 1627.99 | 1579.98 | 1709.00 | 4.26 | 4.23 | 0.12 |
| B10F | 1287 | 1380.52 | 1339.85 | 1452.00 | 4.07 | 4.04 | 0.15 |
| B11F | 2112 | 2279.86 | 2196.84 | 2394.00 | 5.26 | 5.23 | 0.03 |
| B12F | 2035 | 2187.61 | 2122.32 | 2306.00 | 4.76 | 4.73 | 0.06 |
| B13F | 1717 | 1842.93 | 1779.76 | 1933.00 | 4.86 | 4.83 | 0.04 |
| B14F | 1672 | 1853.64 | 1767.82 | 1942.00 | 4.74 | 4.71 | 0.05 |
| B15F | 1789 | 1884.81 | 1832.46 | 1971.00 | 4.96 | 4.93 | 0.05 |
| B16F | 1402 | 1505.79 | 1454.68 | 1577.00 | 4.62 | 4.60 | 0.04 |
| B18F | 1920 | 2046.37 | 1970.35 | 2144.00 | 5.23 | 5.20 | 0.02 |
| B19F | 1792 | 1922.33 | 1853.17 | 2014.00 | 5.12 | 5.09 | 0.02 |
| B20F | 5919 | 6302.47 | 6083.55 | 6649.00 | 6.95 | 6.86 | 0.01 |
| B21F | 1917 | 2097.11 | 2001.17 | 2205.00 | 5.19 | 5.15 | 0.02 |
| B22F | 1063 | 1155.34 | 1112.03 | 1214.00 | 4.93 | 4.89 | 0.02 |
| B23F | 1530 | 1616.45 | 1569.35 | 1696.00 | 4.78 | 4.76 | 0.03 |
| B24F | 1842 | 2022.51 | 1926.18 | 2122.00 | 4.78 | 4.75 | 0.03 |
| B25F | 1659 | 1815.10 | 1743.77 | 1901.00 | 5.04 | 5.02 | 0.02 |
| B29F | 1356 | 1447.33 | 1402.73 | 1520.00 | 4.78 | 4.76 | 0.03 |
| B30F | 1673 | 1844.61 | 1760.67 | 1934.00 | 5.09 | 5.06 | 0.02 |

bottom layer water, number of species and OTUs found, and su
| **Phylogenetic Diversity** | **Good's coverage of library(%)** |
| --- | --- |
| 5300 | 99.05 |
| 2250 | 99.71 |
| 1859 | 99.84 |
| 2247 | 99.78 |
| 2149 | 99.77 |
| 1968 | 99.73 |
| 2802 | 99.70 |
| 2874 | 99.72 |
| 2323 | 99.77 |
| 2297 | 99.70 |
| 2301 | 99.80 |
| 2053 | 99.82 |
| 2532 | 99.76 |
| 2440 | 99.76 |
| 6001 | 99.15 |
| 2588 | 99.67 |
| 1731 | 99.57 |
| 2244 | 99.78 |
| 2530 | 99.69 |
| 2345 | 99.73 |
| 1990 | 99.75 |
| 2354 | 99.71 |

**4.**
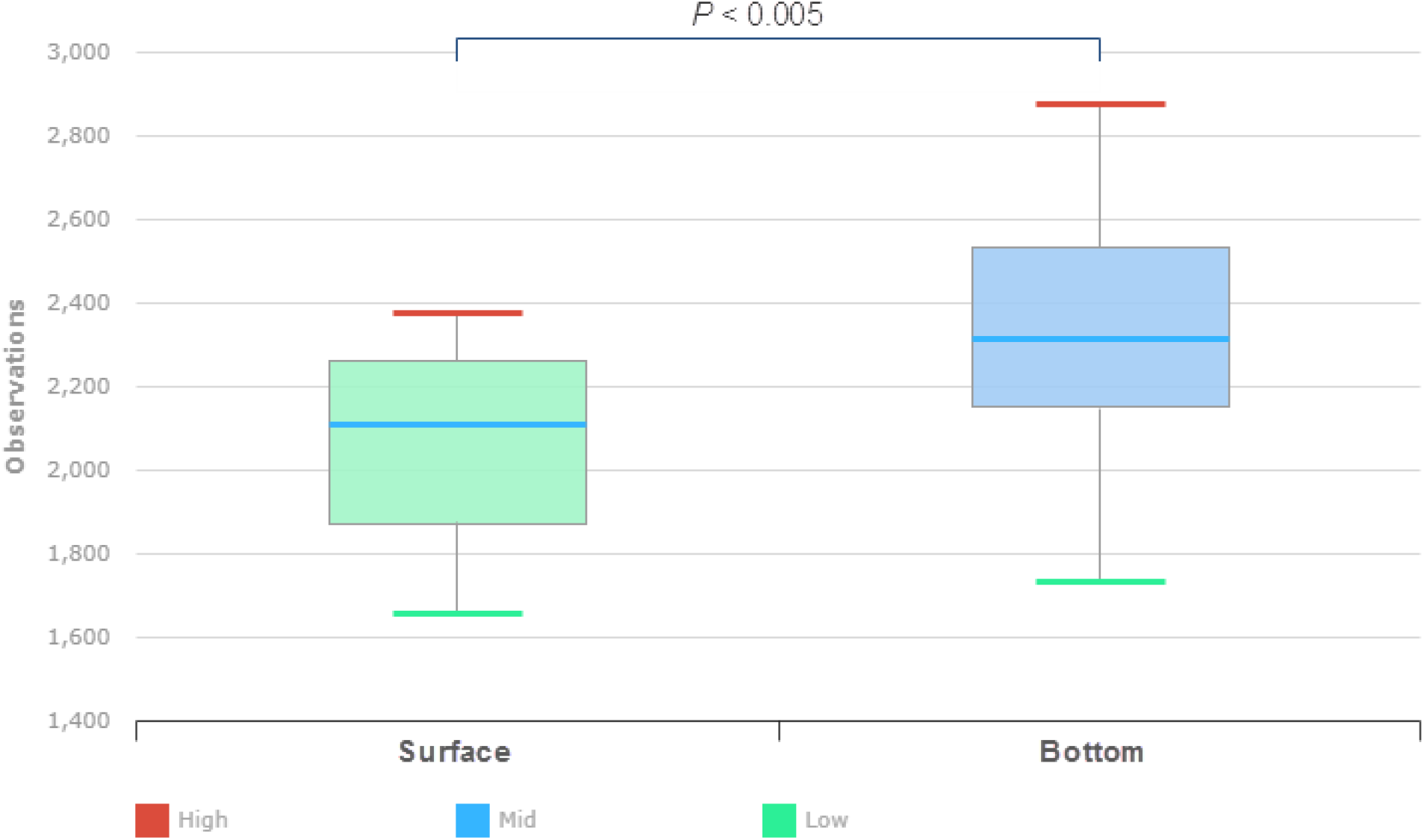
Community diversity represented by phylogenetic diversity index of bacteriome between water samples from the surface and bottom layer.

### 3.3 Identification of Billings microbiome between surface and bottom layer samples

We found seven phyla (*Cyanobacteria, Proteobacteria, Actinobacteria, Bacteriodetes, Verrucomicrobia, Planctomycetes*, and *Chlorobi*) were highly abundant in matched samples from both layers. The phylum *Proteobacteria* and *Bacteroidetes* were less abundant in the surface water samples than in the matched bottom layer samples (**Figure 5**).

**5.**
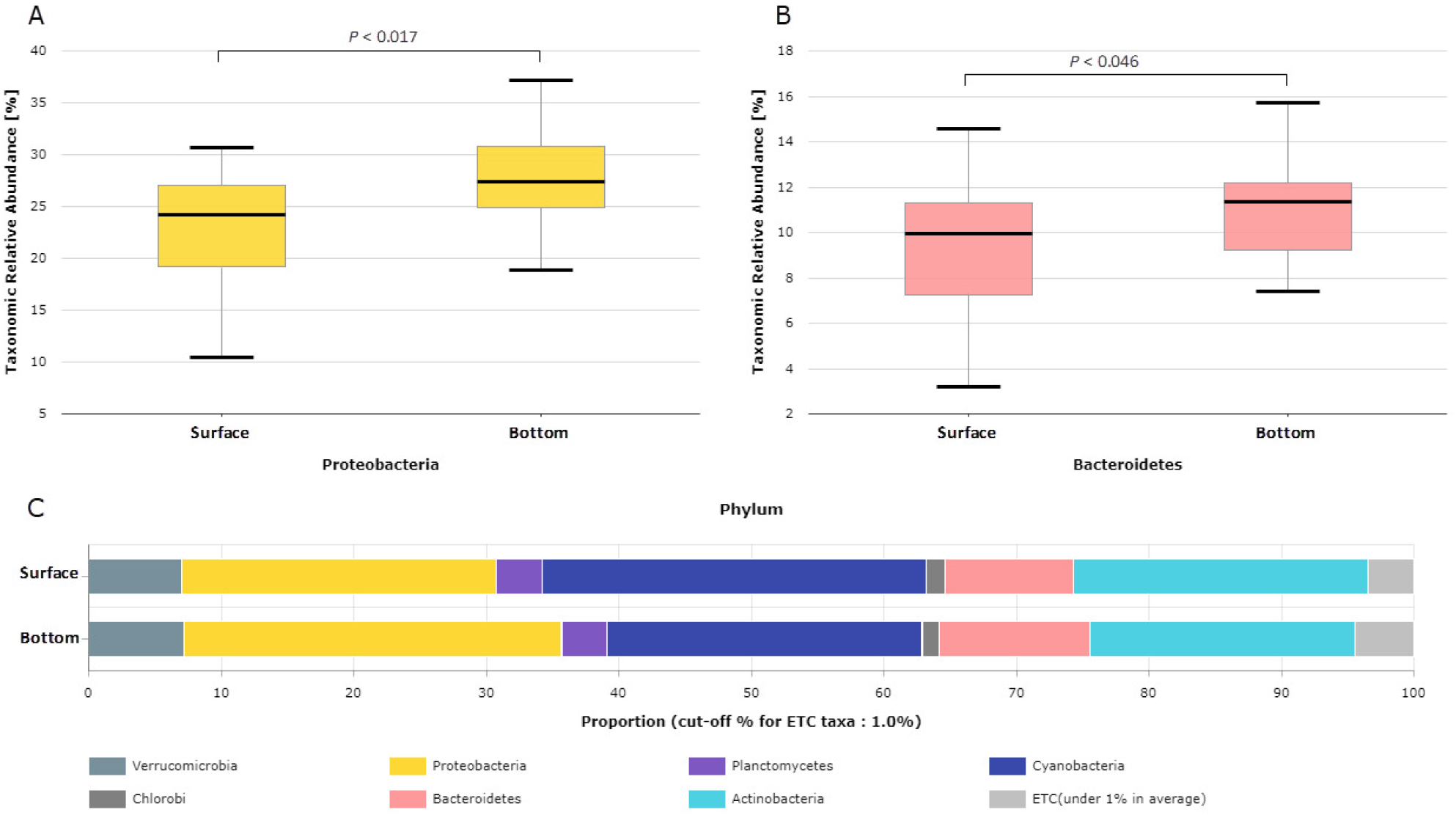
The relative abundance of A) *Proteobacteria* B) *Bacteroidetes*, and C) Phylum level taxonomical abundance of bacteriome between water samples from the surface and bottom layer.

Differentially abundant taxa between the two water samples were identified using the LEfSe algorithm (minimum LDA score: 2.0). This analysis revealed 126 taxa, including 11 class, 24 families, 29 genera, 24 order, 6 phyla, and 32 species. Of these, two families, four genera, one order, and nine species were significantly abundant and discriminative between the groups (FDR-adjusted p-value < 0.02, data not shown). The bacterial species *Stenotrophomonas, Achromobacter, Comamonas, Pseudomonas, Acinetobacter*, and *Schlesneria* were highly abundant in the bottom layer compared to the surface samples (FDR-adjusted p-value < 0.05), while the *Nanopelagicus* species was substantially depleted.

### 3.4 Identification of metabolic-functional pathways between surface and bottom layer

A LEfSe analysis was conducted to identify the most pertinent functional pathways responsible for shaping the Billings microbiome between both layers. We used the software package PICRUSt2 implemented in the EzBiocloud online tool to infer the content of bacterial gene from the data of the16S rRNA gene and aggregated relative abundance of functional genes into metabolic pathways. A total of 83 KEGG orthology (KO) terms were predicted from all OTUs detected in matched samples. Of these, seven differentially abundant (FDR-adjusted P < 0.05) KO terms between surface and bottom layer water were identified (data not shown). Most of the differentially abundant predicted KO terms, including protein metabolism, signaling and cellular processes, and cell motility were highest in bottom layer samples. The pathway of the flagellar assembly was also enriched in the bottom water layer group, which suggests that the growth environment for the bacterial communities in the bottom layer water was much better than that of the surface. PICRUSt module analyses demonstrated that microbiota in the bottom layer exhibits increased biosynthesis of tetrahydrofolate (M00841) and C21-Steroid hormone (M00109) while the surface microbiota showed an increase use of Mce transport system (M00670).

### 3.5 Search for pre-defined bacterial groups and pathogens

The screening for pre-defined bacterial groups in the surface and bottom layer water of Billings reservoir revealed important taxa associated with the human gut that included the phylum *Proteobacteria* (surface; median abundance value 24.7%, bottom layer; median abundance value 27.7%).

The search for the bacterial pathogen in surface water samples based on the criteria described in the Materials and Methods revealed 120 pathogenic genera. Of these, four were potential pathogens for human, animal and plant, 79 for human and animal, one for human and plant, 29 for human, one for animal and plant, two for animal, and 4 pathogenic genera for plant pathogens. Among the 27 surface samples investigated, the genus *Flavobacterium* was the predominant human and animal pathogen, with a median relative abundance of 0.5% and a range of 0.02–1.9%. Of the 31 human pathogens, legionella was the most detected genera with a median abundance of 0.21% and a range of 0.03-0.51%. The search for bacterial pathogens in the 25 bottom layer samples yielded 146, 148, and 18 human, animal, and plant bacterial pathogens, respectively. Again, the genus *Flavobacterium* was the predominant genera (median relative abundance = 0.5% and range of 0.2–3.5%) followed by the plant pathogenic genus *Polynucleobacter* (median relative abundance = 0.9% and range of 0.5–1.8%) and the human pathogenic genus *Stenotrophomonas* (median relative abundance = 0.1% and range of 0.02–0.8%).

### 3.6 Amplification of Cyanotoxin Genes

To investigate the occurrence of potential toxic microcystin-producing strains in the reservoir, total DNA of the microbial community from the water samples that displayed heavy contamination with *Cyanobacteria* were selected and evaluated by conventional PCR assay. The three selected genes DNA that involved in the biosynthesis of *mcy* (*mcyA, mcyE*, and *mcyD*) were successfully amplified in all analyzed samples, confirming the genetic potential of the strains in the reservoir to produce microcystin (data not shown).

## 4. DISCUSSION

In the current study, the relatively high Shannon diversity indices of 2.38–5.39 and 4.04-6.86 and the detection of 305–2207 and 1063-5919 OTUs in surface and bottom layer water, respectively, showing a greater level of overall biodiversity. Perhaps the greatest value of biodiversity is attributed to intensive anthropogenic disturbance in the reservoir that possibly led to increased nutrient discharges, which resulted in higher nitrogen and phosphorous concentrations, as evidenced in this study. It is also possible that other variables that were not included in our study could be contributed to the bacterial community variability in the Billings reservoir. The observed bacterial richness and diversity indices in this study were comparable to those in other reservoirs (21). Taxonomical composition revealed that the *Proteobacteria* phyla were most common, followed by *Cyanobacteria* and *Actinobacteria*. These results are consistent with other Brazilian studies that describe the microbial communities in the Amazon basin (22–24) and Tocantins River (25) but somewhat different from those previously described for reservoirs in China. For instance, Qu J *et al* used the 16S rDNA Illumina approach and showed that *Firmicutes, Proteobacteria, Cyanobacteria*, and *Bacteroidetes* were the dominant bacterial phyla in the Miyun Reservoir, which is considered the largest man-made reservoir in North China. These differences of dominant phyla may be due to a variety of environmental factors including air, soil, and water pollution, and rainfall-induced nutrient fluctuations, and changes in local conditions. Although bacterial richness indices varied little between the surface and bottom layer samples in this study, some specific bacterial groups showed a clear difference. For instance, members of the *β-Proteobacteria* and *γ-proteobacteria, Acidobacteria, Bacteroidetes*, and *Sphingobacteriales* were significantly abundant in the bottom layer than those in the surface water. Since the water flow within a certain layer could contribute significantly to the displacement of bacterial populations, it is reasonable to assume that the latter could constitute a contribution to the observed bacterial community differences between both layers. The difference may also be attributed to the use of distinct DNA extraction methods and/or primers selection (26). It is also conceivable that the phosphorus availability in the bottom layer compared to the surface water could positively influence the growth rates of these bacterial groups (27).

In addition to the seven dominant phyla in Billings’s reservoir, there were approximately 3.5% and 4.4% of unclassified OTU’s labeled as ETC in the surface and bottom layer water, respectively, indicating that as-yet-unidentified bacterial populations with unknown metabolic functions are an important part of the reservoir bacteriome, these findings warrant further analysis.

Members of *Cyanobacteria* were detected as the most dominant genus, with 29 % and 23.7% average abundance in all samples from the surface and bottom layers, respectively, and all *Cyanobacteria* bloom tested here were toxic. These results lend further support to previous studies that demonstrated Brazilian semiarid reservoirs harbor cyanobacterial communities (15, 28–32). The presence of these bacteria in high abundance in the reservoir can be linked to uploading nutrients like ammonia and to the increase in water temperature (20.7°C) at the time of sampling. OTUs of these bacteria are known to out-compete other planktonic microbes for nutrients in eutrophic systems (33). A previous study on the Neuse River, North Carolina, conducted by Paerl recommended that a reduction of 30-40% of NO_3_ had the optimum power of minimizing *M*.*aeruginosa* as a dominant phytoplankter (34). Concerning temperature, there is a consensus among researchers that water temperature below 20^0^C is generally considered unfavorable for the development of the common water bloom forming-genera like *Anabaena* and *Microcystis*. In contrast to this, our finding showed elevated concentrations of *Microcystis* in the reservoir. There are various effects of increasing numbers of *Cyanobacteria* in freshwater systems including oxygen depletion, fish mortalities and toxicity that poses a wide range of health hazards (35). Because the *Cyanobacteria* detected in this study are capable of producing toxin, their presence in the reservoir poses biggest threats to animal and human health.

In this study, *Proteobacteria* represented a second huge portion of the bacterial population in the reservoir, which is similar to that in other studies (36, 37). Despite the dominance of *Proteobacteria* in both layers, a significant difference in the composition of these members was observed at detailed taxonomic analysis. *Betaproteobacteria* were the most detected *Proteobacterial* group in this work, which agrees with other studies (38, 39). OTUs belonging to the *Betaproteobacteria* class have been associated with anthropogenic activities (40). The *Alpha* and *Gammaproteobacteria* found in both layers probably indicate an increase of organic and inorganic inputs and phytoplankton production (27, 38).

The *Actinobacteria* phylum was the third most abundant corresponding to 22.2 and 20% of the reads in the surface and bottom layer samples, respectively. OTUs of this phylum are among the most abundant groups in freshwater habitats (38, 41). Abundances of these microbes inversely correlated with those of *Cyanobacteria* that cause prolonged and irretrievable ecological disturbances to freshwater ecosystems and serve as sentinels of impending ecological damage (42, 43). OTUs of *Actinobacteria* were less abundant when compared to *Proteobacteria*, which comprised 23.7% and 28.5% of the total bacterial abundance in the surface and bottom layer water of Billings reservoir, respectively. *Actinobacteria*, which are well-recognized soil bacteria (44), are frequently detected in oligotrophic freshwater habitats (45), and are often associated with oligotrophic ecosystems (46). They have long been known to produce pigments that protect them against UV radiation, which easily penetrates deep into a freshwater habitat (38).

The alpha diversity analysis conducted between the surface and bottom layer revealed high phylogenetic diversity indices of the bacteriome inhabiting the bottom layer. This result may indicate that bacterial communities in the bottom layer have experienced high diversification rates or immigration of multiple lineages that radiated successfully. One factor that may promote higher bacterial diversification in the bottom layer is that this habitat is possibly less extreme than that of the surface layer and permit easier radiations (47). An examination of Billings water did not reveal significant differences in microbial community beta diversity between the two layers, suggesting that a core microbiome exists between both layers of the reservoir.

Application of the LEfSe method demonstrated the presence of *Stenotrophomonas species* as the most significant specific biomarkers in the bottom layer. These species play an important ecological role in the nitrogen and Sulphur cycles and several *Stenotrophomonas species* can engage in beneficial interactions with plants, promoting growth and protecting plants from attack (48). Since the reservoir contains a huge number of *Cyanobacteria*, the death of many of these bacteria could lead to the release of high content of sulfur-containing amino acids from their cells (49) resulting in sulfur-rich water and thus explain the detection of *Stenotrophomonas species*.

Many countries in the developing world are experiencing intense contamination of their freshwater resources by bacterial pathogens, which have been caused to various waterborne disease outbreaks (50). In the current study, we found that potentially pathogenic bacteria were ubiquitous across all the sampled waters in Billings reservoir. Among the human potential waterborne pathogens, *Legionella* and *Flavobacterium* genera were the most prevalent in both layers; followed by *Stenotrophomonas*. Of note, a study conducted between 2007 and 2009 in the United States by Hlavsa *et al* (51) reported that 21% of the waterborne outbreaks were caused by diverse pathogenic bacteria including *Legionella species*. A report from the river Tama (Tokyo, Japan) showed that the predominant bacteria genus was *Flavobacterium* (Bacteroidia), a freshwater fish pathogen

In addition to taxonomic compositions, we identified a pathway containing genes encoding for flagellar assembly that were present at the greater relative abundance in the Billings metagenome. The abundance of flagellar assembly within the bottom layer indicates that the reservoir contains abundant bacterial communities, which utilize flagellum for its locomotion. It is possible that the adhesion process of these bacteria was influenced by environmental factors like pH and higher levels of metals introduced to the reservoir as a consequence of the rapid expansion of industry and increases in domestic activities

The main limitation of this study is that is that we restricted the analysis to samples at a single point in time. While the spatiotemporal data is not yet complete, analysis of the data already in hand is well underway. In addition, we used bacterial DNA genomics for this investigation, which would have revealed the presence of bacterial populations regardless of whether they are dead or alive, culturable cells, or non-culturable cells.

## 5. Conclusion

In conclusion, this study provides important information about the numerous bacteria inhabiting the Billings reservoir and the combination of environmental that shaped the structure. These results may help pave the way for future studies devoted to control and improve the water quality in the Billings reservoir, which is facing rapid urban development and urbanization.

## Authorship contribution statement

Marta Angela Marcondes: Data curation, Visualization. Investigation, Methodology. Andrezza Nascimento: Investigation, Data curation, Visualization. Rodrigo Pessôa: Formal analysis, Visualization. Jefferson Russo Victor: Writing - review & editing, Visualization. Alberto José da Silva Duarte: Writing - review & editing, Visualization: Sabri Saeed Sanabani: Conceptualization, Supervision, Project administration, funding acquisition.

## Acknowledgements

This work was supported by grant 2018/08631–3 from the Fundação de Amparo à Pesquisa do Estado de São Paulo.

